# Identifying Executive Function Profiles in Children’s Design Fluency: A Clustering Study

**DOI:** 10.1101/2025.08.19.670430

**Authors:** Myriam Sahraoui, Vanessa Hadid, Karim Jerbi, Bruno Gauthier

## Abstract

Executive functions continue to develop throughout childhood and vary considerably across children. Design fluency tasks are commonly used to assess executive functioning through design production, repetitions, and strategy use. When averaged across the full task, these measures summarize performance but do not show how they change as the task progresses. Whether examining these measures minute by minute provides additional information about variability across children remains unclear. We examined this question in 113 neurotypical children aged 7–13 years using a tablet-based design fluency task. To identify performance profiles, we applied K-means clustering separately to design production, repetitions, and strategy use averaged across the task and measured across five one-minute intervals. Overall task measures identified two overlapping profiles. One showed lower design production and lower proportions of repetitions and strategies, whereas the other showed higher production and higher proportions of repetitions and strategies. Minute-by-minute clustering produced nearly identical assignments, but weaker separation, suggesting that temporal information added little to profile identification. However, examining changes over time revealed subtler differences: strategy use followed broadly similar patterns across profiles, but showed a pronounced between-profile difference during the first minute that narrowed thereafter. In contrast, the difference in repetition rates increased as the task progressed, a pattern that may partly reflect differences in overall design production. Profiles did not differ significantly in age, sex, or WISC subtest scores. These findings suggest that overall task measures more clearly distinguish performance profiles, whereas minute-by-minute measures provide complementary information about how these differences evolve over time.

## Introduction

Executive functions (EFs) consist of a set of cognitive processes essential for self-regulation, purposeful behavior, and successful adaptation to new or complex tasks (Diamond, 2013). These cognitive abilities, including inhibition, cognitive flexibility, working memory, and planning (Miyake & Friedman, 2012), continue to develop throughout childhood and adolescence (Best & Miller, 2010) and are especially important during this developmental period as they support critical outcomes spanning academic achievement, social skills, emotional regulation, and long-term cognitive and adaptive functioning (Judd et al., 2024; Kamphorst et al., 2021; Nesayan et al., 2019). Substantial interindividual variability in executive functioning has been observed among typically developing children (Kochhann et al., 2018; Mareva & Holmes, 2024; Muñoz-Parreño et al., 2023). Characterizing this variability could help clarify the different ways children approach EF tasks, potentially informing individualized educational and developmental strategies (Astle et al., 2019; Judd et al., 2024).

EF performance is usually characterized using overall task scores, such as the total number of correct responses produced during a task, interpreted against predefined normative criteria (Astle et al., 2019). While these measures are effective in assessing specific executive components and predicting cognitive outcomes (Bablekou et al., 2023; Chen et al., 2022; Scope et al., 2010), they summarize performance at the level of the whole task and do not capture how it unfolds over the course of the task. Minute-by-minute measures, i.e., measurements taken repeatedly across the task, provide valuable insights into how performance changes during task execution. Within-task variability is often considered as noise rather than as a meaningful source of information, but recent work challenges this assumption, suggesting that such variability may reflect meaningful individual differences (Judd et al., 2024; McKinney et al., 2020). In design fluency, minute-by-minute measures have proven sensitive to differences between clinical and neurotypical groups. For instance, Gauthier et al. (2016) employed Self-Organizing Maps (SOM; Kohonen, 1982), an unsupervised clustering technique, to compare children with and without ADHD. While the total number of designs produced did not differentiate the groups, their time-based production patterns revealed meaningful distinctions. Similarly, fine-grained minute-by-minute measures of design fluency revealed subtle differences in children with Tourette syndrome (Tessier et al., 2022). In typically developing children, minute-by-minute measures have been used to examine how design productivity changes over time (Hurks, 2013). However, whether they reveal performance profiles among neurotypical children, and whether these profiles differ from those obtained with overall task measures, remains unexplored.

Design fluency tasks are well suited to examine both types of measures. In these tasks, children are asked to generate as many novel designs as possible within a limited time period (Fournier et al., 2020; Gauthier et al., 2016; Stievano & Scalisi, 2016). Performance is generally summarized by the number of designs, repetitions, and strategies produced, and these same measures can also be tracked minute by minute. Performance in these tasks recruits multiple executive functions simultaneously: generating novel designs may draw on cognitive flexibility, avoiding repetitions on inhibition and visual working memory, and applying systematic modifications on planning (Stievano & Scalisi, 2016), which makes it difficult to isolate specific constructs (see Suchy et al., 2017). Consistent with this, the number of strategies and unique designs has been associated with verbal fluency and cognitive flexibility, and the percentage of repetitions has been inversely associated with visual working memory (Bolduc, 2023; Stievano & Scalisi, 2016). More strategy production has also been associated with better performance (Goebel et al., 2009; Lezak, 1995). In a tablet-based version of the task, the number of unique designs was associated with performance on established executive function tasks, including the Stroop, Corsi blocks, and the Tower of London (Bolduc, 2023).

Data-driven clustering offers a way to identify performance profiles in the data without relying on predefined categories of participants (Shah et al., 2024). In typically developing children and adolescents, clustering and other person-centered approaches have identified anywhere from two to six EF profiles, depending on sample and methodology (Astle et al., 2019; Bablekou et al., 2023; Litkowski et al., 2020; Muñoz-Parreño et al., 2023; Usai et al., 2018; Younger et al., 2024). Such profiles have been associated with later self-regulation (Brandt et al., 2024). However, these profiles were derived from several tasks, each summarized by a single overall score (e.g., Baez et al., 2020; Brandt et al., 2024; Chaku et al., 2022; Houwen et al., 2019; Martarelli et al., 2018; Sandilos et al., 2019), leaving the evolution of performance within a task unexamined. In Gauthier et al. (2016), the comparison between children with and without ADHD made it possible to test whether minute-by-minute measures captured differences between known groups. In neurotypical children, no such groups are known in advance. Comparing the profiles obtained from overall task measures and minute-by-minute measures therefore provides a way to examine what information minute-by-minute measures add to the characterization of interindividual variability.

The present study compared performance profiles obtained from overall task measures and minute-by-minute measures of a tablet-based design fluency task in neurotypical children. Specifically, we examined (1) whether performance profiles could be identified from overall task measures of designs, repetitions, and strategies, and (2) whether minute-by-minute measures of the same variables would reveal different profiles or provide additional information about this variability. We hypothesized that performance profiles would emerge from overall task measures, and that minute-by-minute measures would reveal characteristics of, or differences between, profiles that are not apparent in overall task measures, particularly in how strategy production and repetitions evolve over time. Finally, we examined whether the profiles differed in age, sex, and WISC subtest scores.

## Methods

### Participants

The data of 113 children (68 girls) aged 7 to 13 years old (M = 10.28, SD = 1.34) were included in this study. Of the participants, 74% were White, 5% were Black, 4% were Hispanic, 3% were Arab, 2% were First Nations, 2% were Asian, and 10% identified as other. The participants’ data came from a previous study in the Laboratoire d’études en neuropsychologie de l’enfance et de l’adolescence (LÉNEA), (see Laniel & Gauthier, 2022). Four cognitive measures were included: the current design fluency task, the Vocabulary, Matrix Reasoning, and Digit Span subtests of the WISC-IV or WISC-V (see Measures section below for details). All neurotypical participants who completed the design fluency task were included in the present study.

To be included in the dataset, the participants had to be schooled in French since the beginning of their education and speak French at home. The exclusion criteria were: (1) a grade under 60% in French on their school report card, (2) repeating a school year, (3) being followed by a speech therapist or a remedial teacher for written language, (4) a history of brain trauma or a neurodevelopmental disorder diagnosis. Participants who scored more than two standard deviations below the mean for their age in reading ability or intellectual quotient (IQ) (as estimated by the subtest Matrix reasoning) were also excluded. Additionally, participants with inattention or hyperactivity and impulsivity scores more than two standard deviations above the mean were excluded (Laniel & Gauthier, 2022).

### Procedure

This study was approved by the comité d’éthique et de la recherche en arts et sciences de l’Université de Montréal (n°: CERAS-2015-16-080-P). All experiments were performed in accordance with the committee guidelines and regulations, as well as with the principles stated in the Declaration of Helsinki.

Participants were recruited in different primary schools and day camps of six different regions in the province of Quebec, Canada. Parents were given a solicitation form, and if they were interested in participating in the study, they were contacted to verify if they were eligible and make an appointment. Participation was on a voluntary basis. Free and informed consent was obtained for all subjects by their legal guardians.

All children completed a battery of cognitive tests (see Laniel and Gauthier (2022) for more details). During testing, the parents were in the waiting room and completed multiple questionnaires. The administration of the AppFLO design fluency task was always performed first in the assessment. For the administration of the WISC subtests, the standard administration procedure was followed.

### Measures

#### AppFLO - Design fluency

The Design fluency task used in this study is a beta version of the tablet-based AppFLO task, developed in the LÉNEA. In this task, five dots are displayed on the screen, arranged in a standardized configuration resembling the face of a die. The size and placement of the dots remain constant across all devices used for assessment. Using their index finger, participants are instructed to draw a design by connecting at least two dots with straight lines. Once a design is completed, they tap an arrow on the right side of the screen to proceed to the next design. The AppFLO application automatically records the following three variables that were used in the present study’s analyses:

- **Number of designs**: All designs produced by the child.
- **Number of repetitions**: All designs that were repeated.
- **Number of strategies**: The number of strategic modifications applied across designs, including additions, subtractions (adding or removing a line relative to the previous design), rotations (turning a previous design), and reflections (mirroring a previous design).

All these metrics were recorded at one-minute intervals throughout the task. One difference between AppFLO and other design fluency tasks is that only the design currently being produced is visible to the participant, which increases the demands on visual working memory (Bolduc, 2023). Figure 1 provides illustrative examples of three designs produced by a participant.

**Figure 1.**
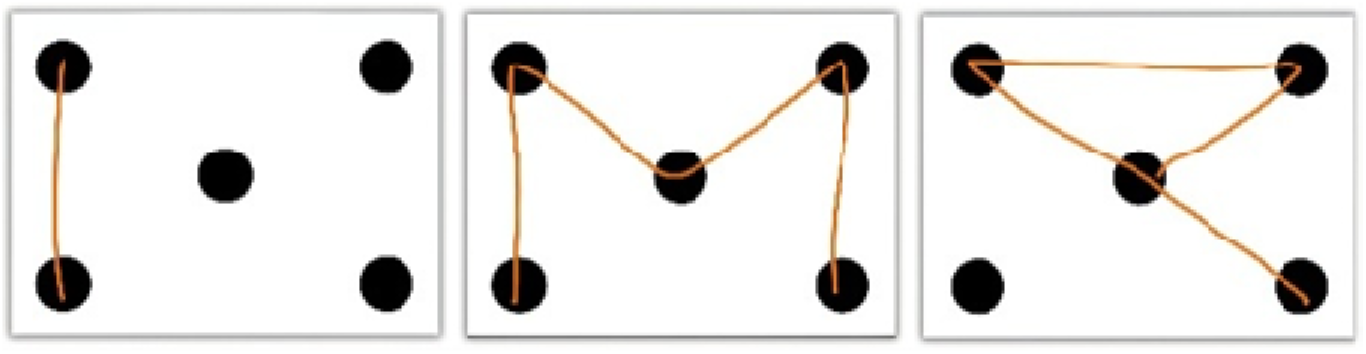
Examples of correct designs completed by a participant on AppFLO.

To account for individual differences in total task productivity, two percentage-based metrics were computed to normalize strategic modifications and repetitions relative to design production. These metrics help reduce the influence of variations in the number of designs produced on the interpretations of strategy production and repetition tendencies:

**(i) Percentage of repetitions**: (Repetitions/Designs) × 100
**(ii) Percentage of strategies**: (Strategies/Designs) × 100

Proportional measures such as these have been widely used in design fluency research to assess executive function performance (Goebel et al., 2009; Stievano & Scalisi, 2016). By normalizing repetitions and strategic modifications relative to designs, percentage-based metrics allow for a fairer comparison across participants and time points.

#### Wechsler Intelligence Scale for Children (WISC)

The WISC-IV (Wechsler, 2005) and the WISC-V (Wechsler, 2015) are standardized measures of intellectual functioning in children aged 6 to 16 years. Depending on the period of data collection, participants completed three subtests from one of the two versions: (1) Vocabulary, consisting of asking the participant to explain the meaning of different words, (2) Matrix Reasoning, consisting of showing participants a series of patterns and asking them to choose an image to complete the missing part of the series, and (3) Digit Span, consisting of asking the participant to repeat increasingly longer sequences of numbers in backward order, or, in the WISC-V only, in ascending order. In the current study, only the backward order score was used, as it was the only working memory-associated metric available for all participants. These subtests have strong validity and reliability (Wechsler, 2005, 2015) and provide measures related to verbal comprehension, perceptual or fluid reasoning, and working memory, respectively.

### Main Analyses

K-means clustering was applied twice: first to the overall task measures, and then to minute-by-minute measures of the task. Overall task measures consisted of the mean number of designs, repetitions, and strategies across the task (3 features). Minute-by-minute measures consisted of the number of designs, repetitions, and strategies at each minute of the task (T1-T5; 15 features). All features were standardized as z-scores before analysis.

To determine whether each dataset showed a tendency to cluster, a Hopkins test was performed before clustering. This test compares the distribution of the data with a randomly generated dataset to evaluate the likelihood of clusters forming. The Hopkins statistic was averaged across 1,000 iterations to reduce variability in the estimate. The H-value varies between 0 and 1, and values above .5 indicate a tendency to cluster (Lawson & Jurs, 1990; Wright, 2022). The Hopkins test was applied to both overall task measures and minute-by-minute measures.

K-means clustering aims to minimize intragroup variance while maximizing intergroup distance. A k-means++ initialization was used to reduce randomness by selecting initial centroids based on a probability distribution. For each representation, solutions from 2 to 10 clusters were tested, and the silhouette coefficient was used to evaluate the quality of the output clusters and to define the optimal number of clusters. The two solutions were then compared using their silhouette coefficients and the agreement between cluster assignments. Minute-by-minute measures were used in two ways: as clustering features, to examine whether they yielded different profiles than overall task measures, and as descriptors of the profiles retained, to examine how differences between profiles unfolded over the course of the task. The K-means clustering, and silhouette coefficients were implemented in Python using the scikit-learn library (Pedregosa et al., 2011).

### Permutation tests

Because the data were not normally distributed, permutation tests were used to compare task performance between clusters. For each comparison, the observed difference in means between the clusters was calculated, followed by 10,000 iterations in which cluster labels were randomly shuffled. A two-sided p-value was computed as the proportion of permutations producing an absolute difference greater than or equal to the observed difference. P-values were corrected for multiple comparisons using the Benjamini-Hochberg False Discovery Rate (BH-FDR) adjustment. Each of the following measures was compared at each time point (T1-T5): (1) the number of designs, (2) the number of repetitions, (3) the number of strategies (additions, subtractions, rotations, reflections), (4) the percentage of repetitions, and (5) the percentage of strategies. Percentages were also compared across the whole task.

Because the retained clusters were derived from the overall task measures (mean number of designs, repetitions, and strategies per minute), differences on these variables are expected by construction. To avoid circularity, these overall task measures were therefore not statistically compared between clusters and are reported descriptively only. Comparisons of the raw measures at individual time points were treated as exploratory analyses describing where in the task between-profile differences were expressed, rather than as independent validation of the clusters. Percentages of repetitions and strategies, which were not directly used for clustering, were also examined exploratorily to determine whether differences in repetitions and strategy use extended beyond differences in overall design production. However, these percentages are not fully independent of the clustering features, as they are computed from the same counts and, given the finite number of possible designs in the task, higher productivity may itself increase the likelihood of repetitions.

### PCA per Cluster

To describe the composition of the clusters identified by K-means, rather than to independently classify or validate them, principal component analyses (PCAs) were run separately within each cluster on the number of designs, repetitions, and strategies at each time point (T1-T5), for a total of 15 features. The PCAs were bootstrapped within each cluster over 1,000 iterations by resampling participants with replacement. For each iteration, components were matched to the original PCA solution based on their similarity, and their signs were aligned to ensure a consistent orientation across iterations. To summarize the contribution of each feature type within each cluster, absolute loadings were summed across time points separately for designs, repetitions, and strategies, with variability estimated from the bootstrap distributions. Because the PCAs were computed independently in each cluster, components are not assumed to represent the same axes across clusters. The analyses were therefore treated as descriptive and exploratory, identifying which measures contributed most to the variance within each profile.

#### Age, Sex, and WISC Subtest Analyses

Sex distribution was compared between clusters using Fisher’s exact test. Mann-Whitney U tests were used to compare age, Vocabulary, Matrix Reasoning, and Digit Span Backward scaled scores between clusters, to examine whether the profiles differed in age and WISC subtest performance. A BH-FDR correction was applied across the Mann-Whitney U tests; Fisher’s exact test for sex was reported separately.

### Correlation analyses

To better examine the relationship between the clusters and age, Spearman rank-order correlations were conducted separately for each cluster, and for each metric of the task. This non-parametric approach was selected for its robustness against non-normal distributions. Correlations were computed using both raw measures (mean number of designs, repetitions, and strategies) and derived metrics (percentage of repetitions and percentage of strategies) collapsed across time points. A BH-FDR correction was performed per cluster across all metrics.

## Results

### Pre-clustering Analysis

A Hopkins statistic was computed to evaluate the clustering tendency of the datasets, averaging 1,000 iterations to account for variability. The Hopkins values obtained were H = .80 for the overall task measures and H = .70 for the minute-by-minute measures.

### Clustering Solution

The optimal number of clusters was determined using the K-means algorithm, with the silhouette coefficient used to assess the separation between clusters (tested across 2 to 10 clusters). The highest silhouette coefficient for the overall task measures (s = .51) indicated a reasonable level of cohesion and separation, supporting a two-cluster solution. Given this result, a two-group clustering solution was retained for further analysis. To visualize the clustering solution, a Principal Component Analysis (PCA) was applied independently to the same dataset used for K-means clustering. The clusters were then projected onto the first two principal components, as shown in Figure 2.

**Figure 2.**
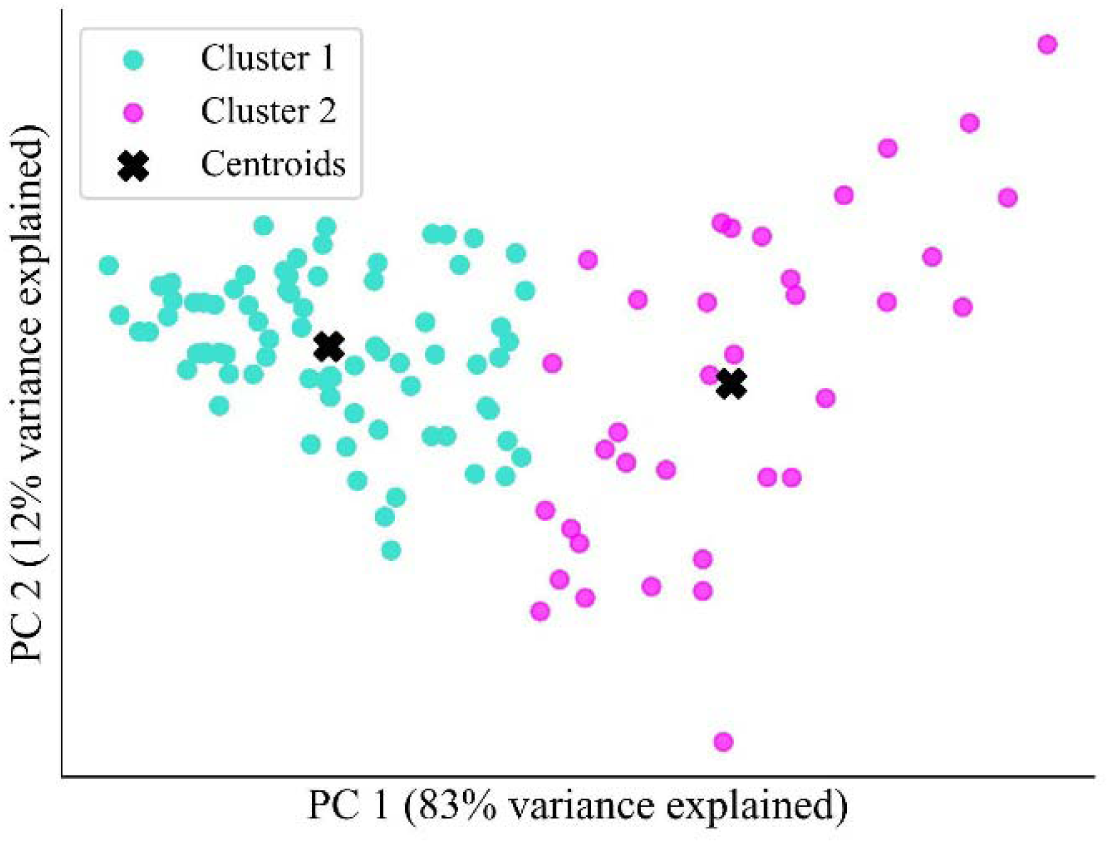
Cluster separation of the K-means solution based on overall task measures, displayed on a PCA. The first two principal components explain **95% of the total variance** and illustrate the separation between the two clusters identified by the K-means algorithm.

The solution based on minute-by-minute measures (15 features) also identified two clusters, but with a lower silhouette coefficient (.33) than the solution based on overall task measures (.51), indicating less clearly separated clusters. Cluster assignments were nearly identical across the two solutions, with only 3 of 113 participants (2.7%) assigned to a different cluster. Although the solution based on minute-by-minute measures (15 features) corresponded to the initially planned analysis, the overall task solution was retained for subsequent analyses given its clearer cluster separation and nearly identical cluster assignments

### Descriptive Characteristics of the Profiles

Table 1 presents the mean and standard deviation of each measure per cluster, along with the mean difference between clusters and exploratory p-values where applicable. Because the retained clusters were derived from the overall task measures, these comparisons describe the profiles rather than validate them. The three overall task measures used to derive the retained clustering solution are presented descriptively without inferential tests. Comparisons at individual time points and for percentage-based measures are exploratory and describe how between-profile differences unfolded over the course of the task.

**Table 1.**
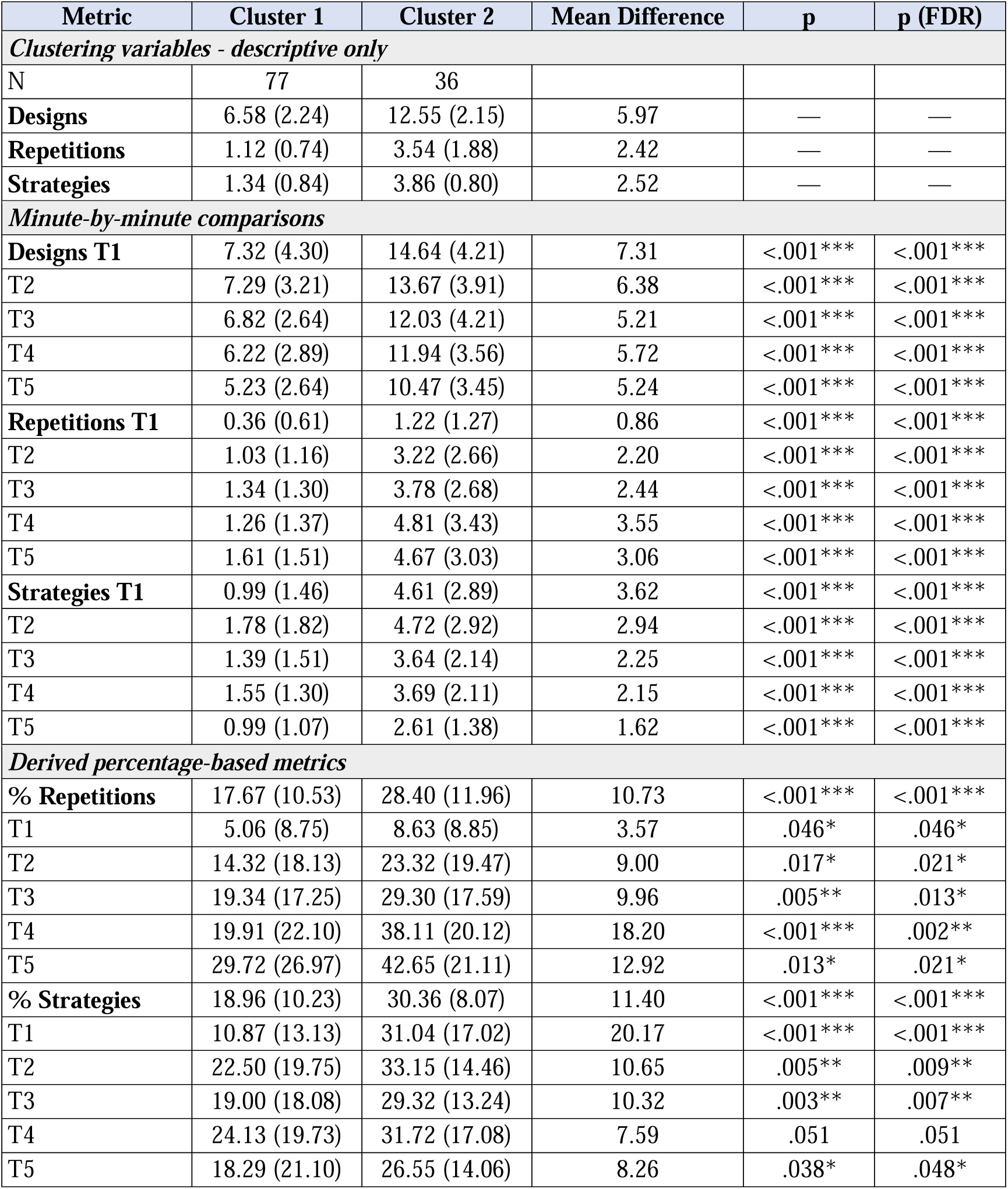
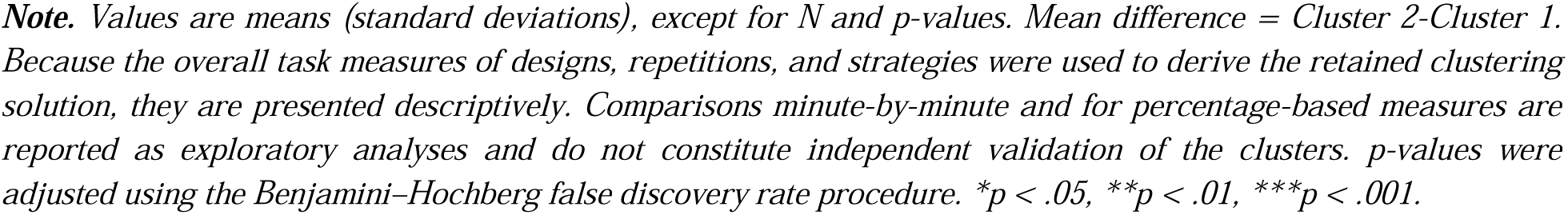
Descriptive characteristics and exploratory comparisons of the two performance profiles.

| Metric | Cluster 1 | Cluster 2 | Mean Difference | p | p (FDR) |
| --- | --- | --- | --- | --- | --- |
| *Clustering variables - descriptive only* |  |  |  |  |  |
| N | 77 | 36 |  |  |  |
| Designs | 6.58 (2.24) | 12.55 (2.15) | 5.97 | — | — |
| Repetitions | 1.12 (0.74) | 3.54 (1.88) | 2.42 | — | — |
| Strategies | 1.34 (0.84) | 3.86 (0.80) | 2.52 | — | — |
| *Minute-by-minute comparisons* |  |  |  |  |  |
| Designs T1 | 7.32 (4.30) | 14.64 (4.21) | 7.31 | <.001*** | <.001*** |
| T2 | 7.29 (3.21) | 13.67 (3.91) | 6.38 | <.001*** | <.001*** |
| T3 | 6.82 (2.64) | 12.03 (4.21) | 5.21 | <.001*** | <.001*** |
| T4 | 6.22 (2.89) | 11.94 (3.56) | 5.72 | <.001*** | <.001*** |
| T5 | 5.23 (2.64) | 10.47 (3.45) | 5.24 | <.001*** | <.001*** |
| Repetitions T1 | 0.36 (0.61) | 1.22 (1.27) | 0.86 | <.001*** | <.001*** |
| T2 | 1.03 (1.16) | 3.22 (2.66) | 2.20 | <.001*** | <.001*** |
| T3 | 1.34 (1.30) | 3.78 (2.68) | 2.44 | <.001*** | <.001*** |
| T4 | 1.26 (1.37) | 4.81 (3.43) | 3.55 | <.001*** | <.001*** |
| T5 | 1.61 (1.51) | 4.67 (3.03) | 3.06 | <.001*** | <.001*** |
| Strategies T1 | 0.99 (1.46) | 4.61 (2.89) | 3.62 | <.001*** | <.001*** |
| T2 | 1.78 (1.82) | 4.72 (2.92) | 2.94 | <.001*** | <.001*** |
| T3 | 1.39 (1.51) | 3.64 (2.14) | 2.25 | <.001*** | <.001*** |
| T4 | 1.55 (1.30) | 3.69 (2.11) | 2.15 | <.001*** | <.001*** |
| T5 | 0.99 (1.07) | 2.61 (1.38) | 1.62 | <.001*** | <.001*** |
| *Derived percentage-based metrics* |  |  |  |  |  |
| % Repetitions | 17.67 (10.53) | 28.40 (11.96) | 10.73 | <.001*** | <.001*** |
| T1 | 5.06 (8.75) | 8.63 (8.85) | 3.57 | .046* | .046* |
| T2 | 14.32 (18.13) | 23.32 (19.47) | 9.00 | .017* | .021* |
| T3 | 19.34 (17.25) | 29.30 (17.59) | 9.96 | .005** | .013* |
| T4 | 19.91 (22.10) | 38.11 (20.12) | 18.20 | <.001*** | .002** |
| T5 | 29.72 (26.97) | 42.65 (21.11) | 12.92 | .013* | .021* |
| % Strategies | 18.96 (10.23) | 30.36 (8.07) | 11.40 | <.001*** | <.001*** |
| T1 | 10.87 (13.13) | 31.04 (17.02) | 20.17 | <.001*** | <.001*** |
| T2 | 22.50 (19.75) | 33.15 (14.46) | 10.65 | .005** | .009** |
| T3 | 19.00 (18.08) | 29.32 (13.24) | 10.32 | .003** | .007** |
| T4 | 24.13 (19.73) | 31.72 (17.08) | 7.59 | .051 | .051 |
| T5 | 18.29 (21.10) | 26.55 (14.06) | 8.26 | .038* | .048* |

**Note.** Values are means (standard deviations), except for N and p-values. Mean difference = Cluster 2-Cluster 1. Because the overall task measures of designs, repetitions, and strategies were used to derive the retained clustering solution, they are presented descriptively. Comparisons minute-by-minute and for percentage-based measures are reported as exploratory analyses and do not constitute independent validation of the clusters. p-values were adjusted using the Benjamini–Hochberg false discovery rate procedure. \* $p < .05$ , \*\* $p < .01$ , \*\*\* $p < .001$ .

#### Number of Designs

On average, participants in Cluster 2 produced more designs than those in Cluster 1 throughout the task (ΔM = 5.97). At T1, Cluster 2 generated 14.64 designs, compared to 7.32 produced by Cluster 1 (ΔM = 7.31, FDR-adjusted p < .001). Both clusters produced fewer designs over time, with Cluster 2 decreasing from 14.64 at T1 to 10.47 at T5, and Cluster 1 from 7.32 to 5.23. The difference between clusters remained present at all time points (ΔM range: 5.21 to 7.31, all FDR-adjusted p < .001; Figure 3).

**Figure 3.**
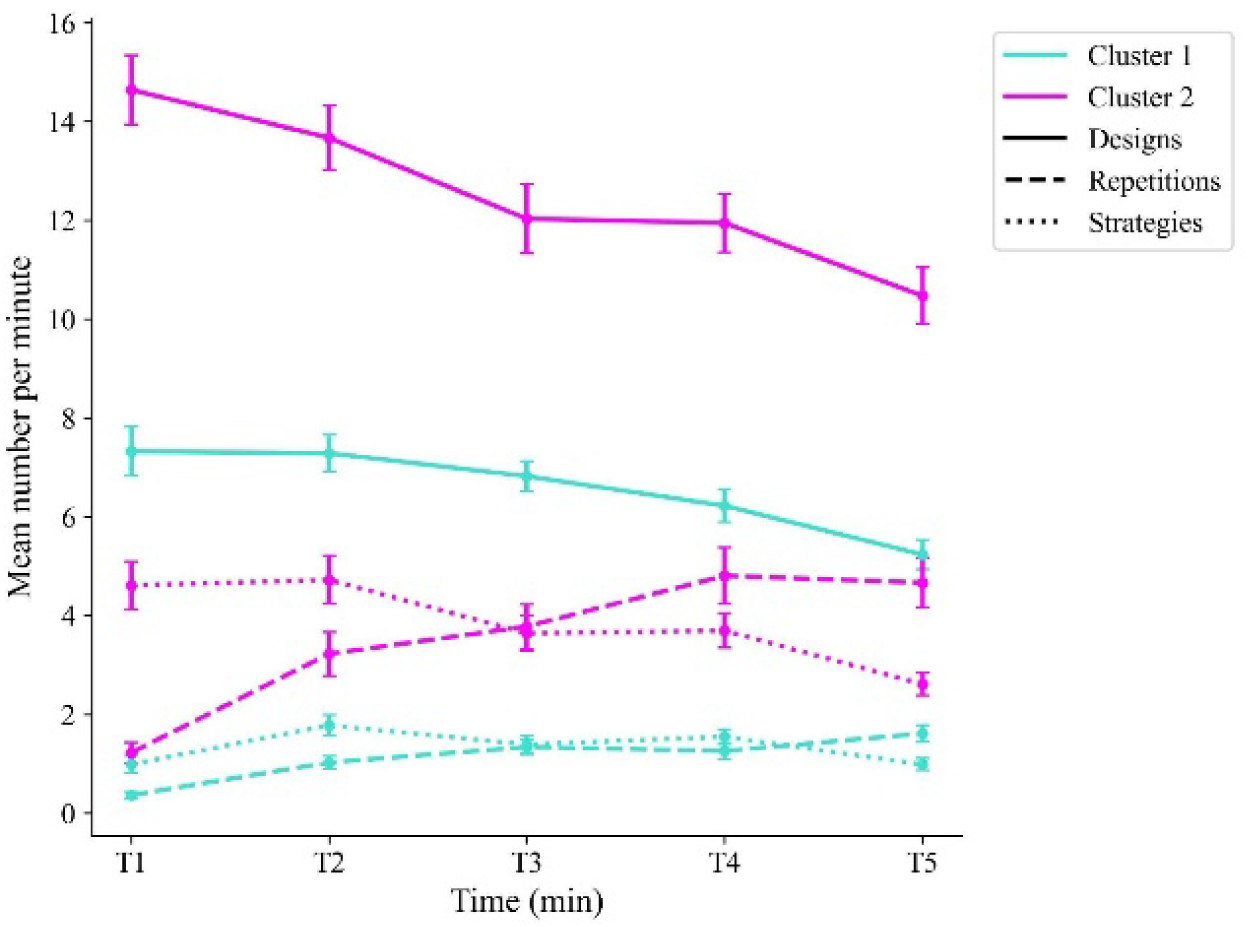
Comparison of the two clusters on the number of designs, repetitions, and strategies across time. Cluster 2 consistently showed higher values than Cluster 1 in designs, repetitions, and strategies. Error bars represent standard error of the mean.

#### Number of Repetitions

On average, participants in Cluster 2 produced about three times as many repetitions as those in Cluster 1 throughout the task (3.54 vs. 1.12; ΔM = 2.42). At T1, Cluster 2 produced 1.22 repetitions, compared to 0.36 in Cluster 1 (ΔM = 0.86, FDR-adjusted p < .001). In Cluster 2, the number of repetitions increased from 1.22 at T1 to 4.67 at T5. Cluster 1 also increased, to a lesser extent, from 0.36 at T1 to 1.61 at T5. The difference between clusters remained present at all time points (ΔM range: 0.86 to 3.55, all FDR-adjusted p < .001; Figure 3).

#### Number of Strategies

On average, participants in Cluster 2 produced about three times as many strategies as those in Cluster 1 throughout the task (3.86 vs. 1.34; ΔM = 2.52). At T1, Cluster 2 produced 4.61 strategies, more than four times the 0.99 produced by Cluster 1 (ΔM = 3.62, FDR-adjusted p < .001). In Cluster 2, the number of strategies decreased from 4.61 at T1 to 2.61 at T5. In Cluster 1, it ranged from 0.99 to 1.78 across time points. The difference between clusters remained present at all time points, and was largest at T1 (ΔM = 3.62) and smallest at T5 (ΔM = 1.62; all FDR-adjusted p < .001; Figure 3)

#### Within-Cluster PCA

For each cluster, a PCA was applied to describe which metrics contributed most to the variance within the profile. For Cluster 1, the first two PCs explained respectively 30.1% and 16.1% of the variance. On the first PC, the number of designs showed the highest loadings, followed by strategies (Figure 4A). On the second PC, designs and strategies at earlier time points (T1-T3) and later time points (T4-T5) loaded in opposite directions. For Cluster 2, the first two PCs explained respectively 31.2% and 13.9% of the variance. On the first PC, the number of repetitions showed the highest loadings, particularly from T2 to T5 (Figure 4B). On the second PC, repetitions at T1 and T2 showed the highest loadings. When loadings were summed by feature type within each cluster (Figure 4C), designs contributed most to the first PC in Cluster 1, followed by strategies and repetitions, while repetitions contributed most in Cluster 2, followed by designs and strategies.

**Figure 4.**
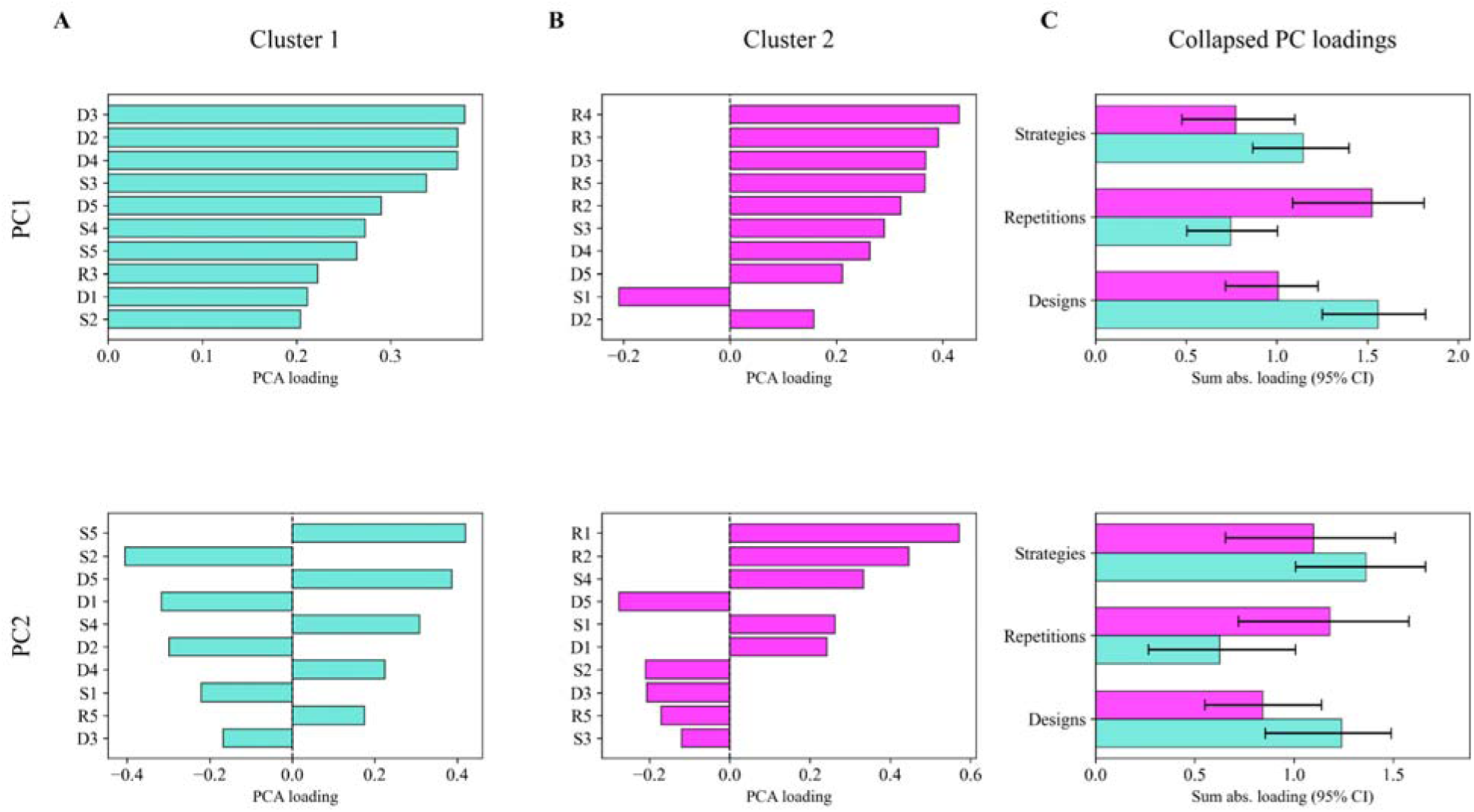
Within-cluster PCA results. Panels A and B show the features with the highest loadings on the first two principal components for Cluster 1 and Cluster 2, respectively. Panel C shows the sum of absolute loadings across time points for each feature type (designs, repetitions, strategies), for both Clusters. Error bars represent 95% confidence intervals across 1,000 bootstrap iterations. D = designs, R = repetitions, S = strategies; numbers indicate the minute (1-5).

### Between-Cluster Comparisons

To account for differences in design production, percentage-based metrics were analyzed, offering a normalized comparison of repetitions and strategies between clusters (Figure 5). While absolute counts of repetitions and strategies provided important insights into group-level differences, percentage scores allowed us to assess patterns relative to overall task performance. However, these percentages remain partially dependent on the number of designs produced.

**Figure 5.**
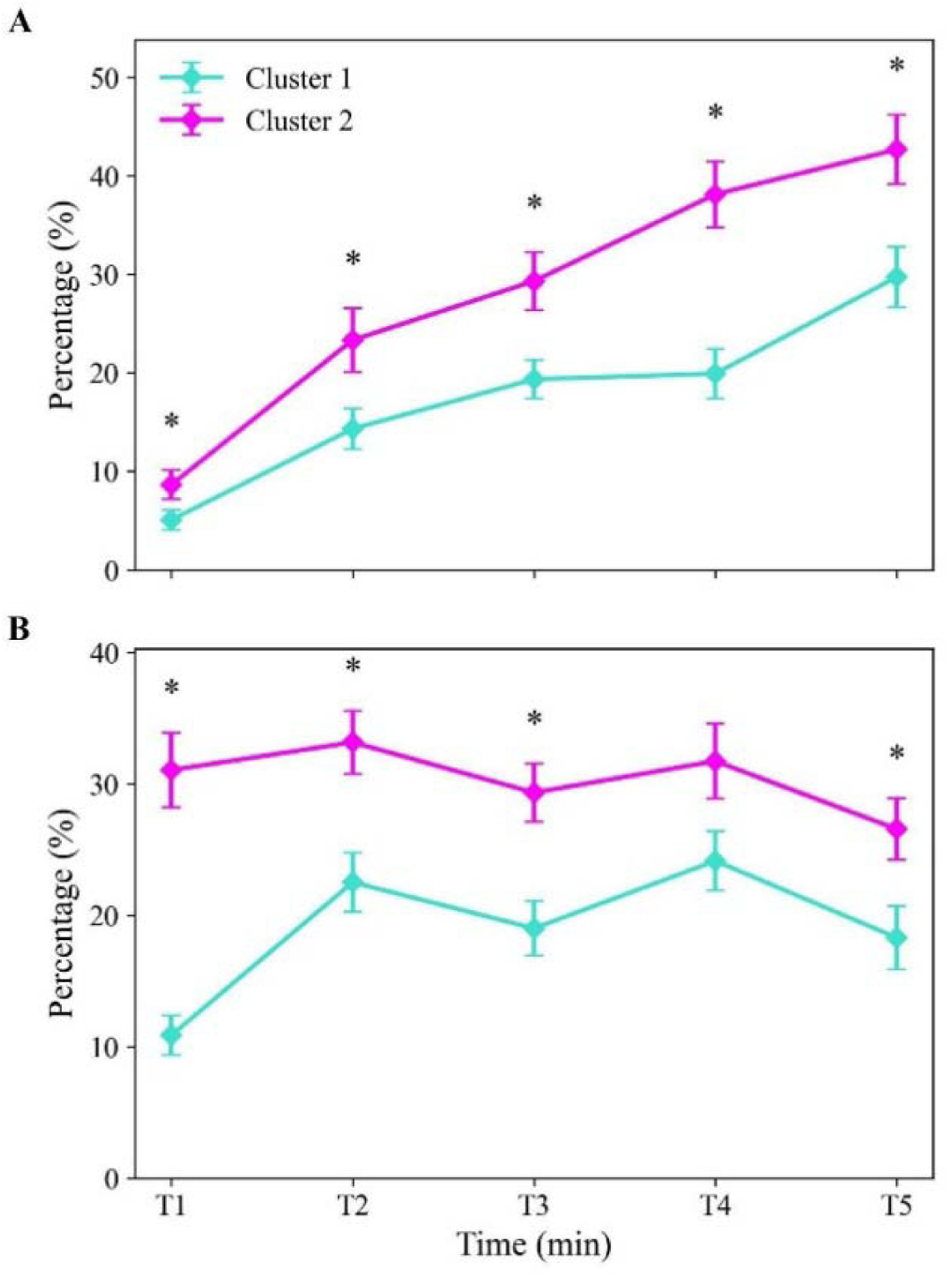
Percentage-based metrics for repetitions and strategies across time points (T1-T5) for Cluster 1 and Cluster 2. Panel A shows the percentage of repetitions and Panel B shows the percentage of strategies. Error bars represent standard error of the mean. *p < .05, BH-FDR corrected.

#### Percentage of Repetitions

Cluster 2 exhibited a significantly higher proportion of repetitions across the task (ΔM = 10.73, FDR-adjusted p < .001), with the most pronounced differences occurring at T4 and T5. At T1, Cluster 2 showed a repetition rate of 8.63%, compared to 5.06% in Cluster 1 (ΔM = 3.57, FDR-adjusted p = .046). This gap generally widened across time. At T5, Cluster 2 exhibited a significantly higher percentage of repetitions, 42.65%, than Cluster 1 (29.72%; ΔM = 12.92, FDR-adjusted p = .021; Figure 5A).

#### Percentage of Strategies

Cluster 2 exhibited overall a significantly higher percentage of strategies (ΔM = 11.40, FDR-adjusted p < .001), with the largest difference at T1. At T1, Cluster 2 used strategies in 31.04% of their designs, compared to 10.87% in Cluster 1 (ΔM = 20.17, FDR-adjusted p < .001). From T2 onward, the difference between clusters was smaller (ΔM range: 7.59 to 10.65) and remained significant at all time points except T4 (ΔM = 7.59, FDR-adjusted p = .051; Figure 5B).

#### Age, Sex, and WISC Subtest Scores

Mann-Whitney U tests revealed no significant differences between clusters in age (Cluster 1: M = 122.29 months; Cluster 2: M = 125.56 months; U = 1201.5, FDR-adjusted p = .513) or in WISC scaled scores, including Vocabulary (U = 1172.0, FDR-adjusted p = .513), Matrix Reasoning (U = 1370.0, FDR-adjusted p = .993), and Digit Span Backward (U = 1358.0, FDR-adjusted p = .993; Table 2). Sex distribution did not differ significantly between clusters either (Cluster 1: 62.3% female; Cluster 2: 55.6% female; Fisher’s exact test, p = .540). The same comparisons conducted on the solution based on minute-by-minute measures (15 features) also revealed no significant differences between clusters in age, sex, or WISC scaled scores.

**Table 2.** Comparison of sex, age, and WISC subtest scaled scores between clusters.

| **Metric** | **Cluster 1** | **Cluster 2** | **U** | **p** | **p (FDR)** |
| --- | --- | --- | --- | --- | --- |
| **Sex (% female)** | 62.3% | 55.6% |  | .540 |  |
| **Age (months)** | 122.29 (16.54) | 125.56 (15.01) | 1201.5 | .257 | .513 |
| **Vocabulary** | 10.78 (2.51) | 11.47 (2.98) | 1172.0 | .220 | .513 |
| **Matrix Reasoning** | 10.64 (3.04) | 10.61 (2.63) | 1370.0 | .993 | .993 |
| **Digit Span Backward** | 9.74 (2.56) | 9.71 (2.6) | 1358.0 | .860 | .993 |

**Note.** Values are means (standard deviations), except for sex. Sex was compared using Fisher's exact test; age and WISC subtests were compared using Mann-Whitney U tests, with p-values corrected using the Benjamini-Hochberg FDR procedure. \* $p < .05$ , \*\* $p < .01$ , \*\*\* $p < .001$ .

#### Age-Related Correlations

Spearman correlations were conducted separately for each cluster to examine the relations between age and each design fluency metric (number of designs, repetitions, and strategies, and percentages of repetitions and strategies), collapsed across time points. No significant correlations were found.

## Discussion

Overall task measures are commonly used to characterize EF interindividual variability in typically developing children, but they summarize performance at the level of the whole task and do not capture how it unfolds over time. This study therefore examined (1) whether performance profiles could be identified from overall task measures of a design fluency task, and (2) whether minute-by-minute measures would reveal different profiles or provide additional information about this variability. Using an unsupervised clustering method, profiles were identified in the data. The results revealed two overlapping performance profiles, differing not only in the number of designs produced but also in the percentages of strategies and repetitions.

Cluster 1 showed a lower-output profile, with lower fluency throughout the task (fewer designs) and a slower increase in repetition errors (percentage of repetitions) over time. While the percentage of strategies remained lower than those of Cluster 2, they increased after T1 compared to their own early performance. Furthermore, additional within-cluster PCA suggested that the production of designs contributed the most to the variance in Cluster 1, compared to repetitions and strategies.

Cluster 2 exhibited a high-output profile throughout the task, generating more designs but with a steeper increase in repetition errors (percentage of repetitions) and a higher percentage of strategies, particularly at T1. The within-cluster PCA further characterized these profiles: design production contributed most strongly to the first principal component in Cluster 1, whereas repetitions contributed most strongly in Cluster 2.

To complement the comparison between overall task and minute-by-minute measures, within-cluster PCA was used to further characterize each profile by identifying which features of minute-by-minute performance contributed most to variability within that profile. Design production contributed most strongly to the first principal component in Cluster 1, whereas repetitions contributed most strongly in Cluster 2. Thus, although minute-by-minute measures did not substantially alter profile assignment, they provided additional information about the dimensions of performance variability within each profile.

Clustering based on minute-by-minute measures yielded nearly identical cluster assignments (only 3 of 113 children changed cluster), but a weaker separation than clustering based on overall task measures. The Hopkins statistic indicated a tendency toward clustering for both overall task (H = .80) and minute-by-minute measures (H = .70). For overall task measures, the silhouette coefficient (.51) corresponded to a reasonable separation between clusters (Kaufman & Rousseeuw, 1990, p. 88), compared to .33 for minute-by-minute measures. Although both representations were derived from the same performance data, the near-identical grouping indicates that similar participant assignments were recovered across the two representations. However, the clusters were not fully distinct: several children were located near the boundary between clusters (Figure 2), and the large within-cluster variability (Table 1) indicates that the distributions of the two clusters overlapped on most measures. These results are therefore best described as two overlapping profiles rather than discrete categories. Conceptualizing performance variability as behavioral profiles may help better understand the different ways children approach this type of task and how these approaches may influence performance.

Rather than competing, overall task and minute-by-minute measures appeared complementary: overall task measures yielded more clearly separated profiles, while minute-by-minute measures described how the characteristics distinguishing these profiles unfolded over the course of the task. Some temporal patterns were similar between clusters, with both producing fewer designs over time, as previously reported for the Five-Point Test (see Woods et al., 2016), and both showing an increase in the percentage of repetitions. In particular, the between-profile difference in strategy use was greatest during the first minute and narrowed thereafter, whereas the difference in repetition rates generally increased as the task progressed. Our hypothesis was therefore partially supported: minute-by-minute measures did not lead to different profiles, but they revealed how differences between profiles evolved over the task. These findings suggest that differences between profiles were not captured by design production alone but also involved the relative occurrence of repetitions and strategies.

### Cluster Differences in Output-Repetition Tradeoffs

A central descriptive finding was the co-occurrence of higher design production and a higher proportion of repetitions in Cluster 2, particularly as the task progressed. We refer to this pattern as an output-repetition tradeoff. This notion is conceptually related to the speed-accuracy tradeoff, which refers to the well-established principle that more accurate responses typically require more time, necessitating a balance between speed and accuracy that can vary across both individuals and situations (Heitz, 2014). However, as response times were not measured in the present study, we refer to a tradeoff between output and repetitions rather than between speed and accuracy. This notion also echoes the error ratio proposed for the Ruff Figural Fluency Test, which relates the number of perseverative errors to the number of unique designs to capture the ability to maximize production while minimizing repetitions (Ross, 2014; Ruff, 1996). Cluster 2 consistently generated more designs than Cluster 1 and showed a larger increase in relative repetitions over time, whereas Cluster 1 produced fewer designs and showed a more modest increase in repetitions. Importantly, this pattern does not establish that children deliberately prioritized output, monitoring, or caution. In contrast, Cluster 1 produced fewer designs but showed a smaller increase in repetition errors, which may reflect a focus towards monitoring their repetitions over global production.

These divergent patterns suggest that Cluster 2 may have prioritized fluency, potentially at the expense of monitoring their responses. This interpretation is consistent with the higher percentage of repetitions in Cluster 2 and the importance of repetitions in the variance of this profile. In Cluster 1, while producing less, the number of designs contributed the most to the composition of this profile. This pattern may reflect a more cautious approach, in line with models emphasizing the evolving interplay between fluency, flexibility, planning abilities and inhibition (Dumont et al., 2022; Goodgold-Edwards et al., 1997; Rommelse et al., 2016).

The inhibitory resource depletion hypothesis offers a possible additional explanation for this pattern, suggesting that sustained cognitive effort gradually depletes attentional and inhibitory resources, increasing reliance on automatic responses and, consequently, repetition errors (Dumont et al., 2022; Poreh, 2022). This could be consistent with the rise in repetition errors in Cluster 2, whose percentage of repetitions was significantly higher than that of Cluster 1 at all time points, with differences generally increasing across time. In contrast, the more modest rise in repetition errors in Cluster 1 may reflect that maintaining a lower output helped limit repetitions over time, although it may also partly reflect fewer opportunities to repeat a design when fewer designs are produced. Working memory demands imposed by the task format may also contribute to the difference between the clusters. Unlike traditional fluency tasks, where participants can visually track their previous designs, the tablet-based version displayed one design at a time, requiring children to rely more heavily on working memory to monitor their responses and avoid repetitions. Because Cluster 2 consistently generated more designs than Cluster 1, they had more past responses to monitor, potentially increasing the load on working memory resources. However, scores on the Digit Span Backward subtest did not differ significantly between clusters, so the observed differences in task performance cannot be directly attributed to this specific measure of working memory. As fatigue, resource depletion, and working memory load during the task were not directly measured, these interpretations remain speculative.

### Age, Sex, and General Cognitive Abilities

In the current study, age, sex, and the WISC subtest scores did not differ significantly between clusters, for both overall task minute-by-minute clustering solution, and no significant correlation with age was found in either cluster. Although studies generally report associations between age and EF task performances (Best et al., 2009; Klenberg et al., 2001), findings remain mixed regarding cognitive profiles and design fluency. For example, Regard et al. (1982) found a positive association between age and design fluency, Gabrig et al. (2018) reported significant differences between EF profiles and age, and Brandt et al. (2024) found that age, sex, and processing speed were associated with EF profile membership in a large sample of children aged 6 to 11. However, other studies have found only weak or no correlation between age and performance, or age and cluster separation (e.g., Astle et al., 2019; Hurks, 2013; Stievano & Scalisi, 2016). Similarly, some studies report a link between IQ and design fluency (e.g., Goebel et al., 2009; Regard et al., 1982), while others find no significant effects (Hurks, 2013; Tucha et al., 2012). The literature is more consistent concerning sex, with most studies reporting negligible or no effects of sex on EF and design fluency tasks (Albert et al., 2010; Bolduc, 2023; Klenberg et al., 2001). Taken together, these results suggest that the profiles may not be primarily explained by age, sex, or general cognitive abilities. Other factors, such as how children regulate their performance during the task, could contribute to these differences, although this was not directly assessed. However, non-significant results do not rule out smaller effects that the present sample, particularly Cluster 2, may not have been able to detect. Taken together, we did not detect evidence that the profiles differed according to age, sex, or the WISC subtest scores examined here.

### Limitations and Future Directions

Several limitations must be acknowledged. First, although the sample size was sufficient for clustering, it remains relatively modest (n = 113) and predominantly constituted of French-speaking White children from Quebec. Furthermore, the large age range (7 to 13 years old) might have obscured finer changes, as EF development does not follow a strictly linear increase across time. In addition, profiles were identified at a single time point, although longitudinal work suggests that a substantial proportion of children change EF profiles across middle childhood and early adolescence (Brandt et al., 2025). Whether the profiles identified here remain stable over time therefore remains to be examined.

Second, the features used for clustering (designs, repetitions, and strategies) are inherently interrelated parts of design fluency performance. Consequently, the identified profiles should be interpreted as multivariate behavioral patterns reflecting the joint contribution of these measures, rather than the independent influence of any single feature. Moreover, percentages of repetitions and strategies remain partially dependent on overall productivity, as children producing more designs also have more opportunities to repeat a design or use a strategy. The number of repetitions and strategies also showed right-skewed distributions, as previously reported for perseverative errors (Izaks et al., 2011), which may have limited the sensitivity of between-cluster comparisons and may inflate the Hopkins statistic independently of the presence of distinct clusters (Lawson & Jurs, 1990). In addition, as the number of possible designs is limited, children may progressively exhaust the available options, which could impose a similar declining trajectory across children and limit between-profile differences in temporal dynamics.

Third, few measures were available to externally validate or interpret the profiles. Performance variability across the task may have reflected cognitive fatigue, task engagement, working memory load, or processing speed, which may account for a substantial part of individual differences in EF tasks (Löffler et al., 2024) and has been associated with EF profile membership in children (Brandt et al., 2024). As the clusters differed mainly in productivity, future studies should examine whether these profiles persist after accounting for processing speed. Neural measures recorded during the task, such as mobile electroencephalography, could also help distinguish between these processes. Moreover, general cognitive abilities were assessed with either the WISC-IV or the WISC-V, depending on the period of data collection. While relatively few changes were made, this still may have introduced additional variability in these scores.

Finally, the profiles identified here need to be replicated in larger and more diverse samples, and with other methods, including dimensional approaches that may better capture gradual and intermediate differences between children. Finer temporal measures, such as the timing of each individual design, could also help determine whether meaningful temporal differences exist at a smaller scale.

## Conclusion

This study identified two performance profiles in neurotypical children using an unsupervised clustering approach, which may reflect different output-repetition tradeoffs. Cluster 1 showed a lower-output pattern with a more modest increase in repetitions over time, while Cluster 2 exhibited higher fluency but a steeper increase in repetitions. Overall task and minute-by-minute measures revealed broadly similar profiles: overall task measures yielded more clearly separated clusters, while minute-by-minute measures described how differences between profiles unfolded over the course of the task. Beyond the number of designs produced, the relative proportion of repetitions and strategies provided additional information on children’s performance, suggesting that EF assessment may benefit from looking beyond a single measure of output.

Further research is needed to determine whether these profiles remain stable over time or shift with context, experience, or cognitive training, and whether they generalize to larger and more diverse samples. If confirmed, these profiles could contribute to perspectives on cognitive variability in neurotypical populations and to a better understanding of the mechanisms underlying executive control. In the longer term, identifying such profiles could help inform cognitive interventions, for example by supporting children in the high-output profile in monitoring their responses, and children in the lower-output profile in developing their flexibility and fluency. Understanding how these performance profiles relate to learning and problem-solving could also eventually help guide educational strategies.

## Acknowledgments

We would like to thank Sina Esmaeili and Fateme Aliyari for their invaluable support and careful proofreading of this paper.

## Data availability statement

The data and material used in this study are not publicly accessible due to the ethics approval. Data sharing by the authors will be considered upon reasonable request, and within the constraints of the ethical committee approval. The analytic code necessary to reproduce the analyses presented in this paper can be made available upon request to the first author.

## Declaration of generative AI use

Generative AI was used to assist with language editing, proofreading, and restructuring passages based on the authors’ content and analyses. The authors reviewed, edited, and verified all AI-generated suggestions and take full responsibility for the manuscript.

## Author Contribution Statement

B.G., M.S., and K.J. conceived and designed the study. Data were collected in B.G.’s laboratory. M.S. performed the primary data processing and analyses, while V.H. conducted additional analyses and validation. M.S. drafted the initial manuscript. All authors contributed to reviewing and editing the final version and approved it for submission. B.G. and K.J. provided supervision and contributed to funding acquisition.

